# Differential effects of obesity and diabetes on the action potential waveform and inward currents in rat ventricular myocytes

**DOI:** 10.1101/2024.09.03.610949

**Authors:** Anatoliy Shmygol, Gilles Bru-Mercier, Ahmed S. Sultan, Frank C. Howarth

**Affiliations:** College of Medicine and Health Sciences, Department of Physiology, United Arab Emirates University, Al Ain, UAE

**Keywords:** action potential, voltage-gated Na^+^ current, voltage-gated Ca^2+^ current, type 2 diabetes, Zucker rat

## Abstract

Obesity is a significant health concern worldwide, increasing the risk for type 2 diabetes mellitus (T2DM) and cardiovascular disease. Studies have found various vascular anomalies, abnormal heart rhythm, and impaired electro-mechanical coupling in patients with diabetes. Research on non-diabetic obese individuals has shown that besides diabetes-related complications, obesity itself raises the risk of developing cardiovascular disease. Recent studies have revealed a decrease in the speed of electrical signal conduction in the heart, along with slight gap junction dysfunction, which is insufficient to explain the observed impediment of impulse conduction. It’s still unclear whether this impairment is due to obesity-related fat toxicity or diabetes-related factors. Our study aimed to investigate the ventricular action potential parameters and voltage-gated Na^+^ (I_Na_) and Ca^2+^ (I_(Ca, L)_) currents in Zucker fatty (ZF) rats in comparison to Zucker diabetic fatty (ZDF) rats, a well-established model of obesity and T2DM.

Ventricular myocytes were enzymatically isolated from 25-30-week-old Zucker rats. Resting and action potentials were recorded from isolated left ventricular myocytes using a β-escin perforated patch clamp in current-clamp mode; I_Na_ and I_(Ca, L)_ were recorded using whole-cell patch clamp techniques.

Ventricular myocytes from ZF rats showed higher excitability and faster upstroke velocity. ZF rats also had a larger density of I_Na_. Conversely, ZDF rats had decreased I_Na_ which correlated with a reduced velocity of the action potential upstroke. There were no changes in the density or voltage sensitivity of I_(Ca, L)_ among the three groups of animals.

In conclusion, obesity alone and obesity accompanied by DM have distinct effects on the action potential waveform, I_Na_ density and excitability of ventricular myocytes in a rat model of T2DM.

## Introduction

Diabetes mellitus (DM) is the seventh leading cause of death worldwide. In 2017, an estimated 451 million adults aged 18-99 years had DM, which is expected to increase to nearly 700 million by 2045 (1). Obesity, on the other hand, has almost tripled since 1975. Obese people are more likely to develop metabolic syndrome, insulin resistance and type 2 diabetes mellitus (T2DM) (2). The term “diabesity” was created, due to the strong link between obesity and diabetes. However, although most individuals with T2DM are obese, only a fraction of obese individuals develop T2DM (3). Studies in non-diabetic obese patients show that, in addition to diabetes-related complications, obesity itself elevates the risk of developing cardiovascular disease (4).

Cardiovascular disease is a leading cause of mortality in individuals with T2DM. Previous studies have reported a multitude of vascular abnormalities, aberrant heart rhythm and defective electro-mechanical coupling in patients with diabetes. Ethical constraints and practical limitations of studying T2DM in humans necessitate the use of animal models to unravel this disease’s pathogenesis and pathophysiological mechanisms of its complications. Although animal models cannot recapitulate all aspects of human disease, the Zucker diabetic fatty (ZDF) rat has attracted considerable interest as one of the closest to the human condition. The original Zucker fatty (ZF) rat was found to harbour a spontaneous missense mutation designated “fatty” (fa) in the leptin receptor gene. Homozygous animals with this mutation (fa/fa) develop hyperphagia, severe hyperlipidaemia and obesity without hyperglycaemia, while heterozygous (fa/+) animals do not show hyperphagia and remain lean (ZL rats) (5). Subsequent observations revealed that, on rare occasions, the ZF male rats developed severe hyperglycaemia and high insulin resistance. Inbreeding these animals for 10 generations resulted in the development of a strain of fatty rats consistently demonstrating a severe hyperglycaemic phenotype (6,7). The resulting three strains of Zucker rats, namely ZL, ZF and ZDF, provide a convenient model for investigating the cardiovascular effects of obesity separately from those caused by a combination of obesity and T2DM (8–10). Like humans, type 1 and type 2 diabetic animals often show electrical remodelling of the myocardium manifested as long QT interval, impaired contractility and abnormal conduction velocity (9,11–15). Diabetes-induced electrical remodelling is largely attributed to a prolongation of the ventricular action potentials via a decrease in outward K^+^ currents and possible upregulation of I_Ca_ and aberrant conduction of action potentials (12,16–19). The latter is thought to be at least in part due to a malfunctioning of the gap junctions between neighbouring myocytes within the myocardium resulting from an imbalance between phosphorylated and dephosphorylated connexin 43 (Cx43) and/or redistribution of Cx43 (20,21). Olsen *et al*. have described impaired conduction velocity in the myocardium of ZDF rats accompanied by an increased lateralisation of Cx43 (22). The authors concluded that the observed lateralisation was not the sole cause for impaired conduction but additional mechanisms were at play. An important determinant of cardiac impulse propagation is the large and rapidly activating sodium current (23,24). In this paper, we tested the hypothesis that obesity and diabetes modulate ventricular action potential differentially via selective enhancement or reduction of fast sodium current. We found that, in diabetic animals, the rate of rise of the ventricular action potential is substantially reduced in comparison to that in fatty non-diabetic animals. This was mirrored by a substantially reduced I_Na_ implying a causative relationship. The action potentials were of longer duration in fatty and fatty diabetic groups compared to age-matched lean animals.

## Materials and Methods

### Animals

Experiments were performed in 16 ZDF, 16 ZF and 13 ZL male rats (Charles River Laboratories, Margate, Kent, UK) and experiments commenced at 195 days of age. Rats were maintained under a 12-hour light/12-hour dark cycle with free access to water and a standard chow diet. Body weight, heart weight and non-fasting blood glucose (OneTouch Ultra 2, LifeScan) were measured immediately before each cell isolation. Insulin concentration in blood plasma was measured using a commercially available rat insulin ELISA kit (10-1250-01, Mercodia rat Insulin ELISA). Ethical approval for this study was obtained from the Animal Ethics Committee, College of Medicine & Health Sciences, United Arab Emirates University.

### Isolation of ventricular myocytes

Left ventricular myocytes were isolated from the rats according to previously described techniques (25). In brief, the animals were euthanized using a guillotine and hearts were removed rapidly and mounted for retrograde perfusion according to the Langendorff method. Hearts were perfused at a constant flow of 8 ml/g heart/min and at 36–37°C with cell isolation solution containing in mmol/l: 130 NaCl, 5.4 KCl, 1.4 MgCl_2_, 0.75 CaCl_2_, 0.4 NaH_2_PO_4_, 5.0 HEPES, 10 glucose, 20 taurine, and 10 creatine (pH 7.3). Perfusion flow rate was adjusted to allow for differences in heart weight between animals. When the heart had stabilized, perfusion was continued for 4 min with Ca^2+^-free cell isolation solution containing 0.1 mmol/l EGTA, and then for 6 min with cell isolation solution containing 0.05 mmol/l Ca^2+^, 0.75 mg/ml collagenase Type 1 (Worthington Biochemical Corp, Lakewood, NJ, USA) and 0.075 mg/ml type XIV protease (Sigma, Taufkirchen, Germany). Left ventricle tissue was excised from the heart, minced, and gently shaken for 4 min at 36-37 °C in collagenase-containing isolation solution supplemented with 1 mg/ml fatty acid-free bovine serum albumin. The cell suspension was then filtered using 300-micron nylon mesh (Cadisch Precision Meshes, UK). The filtrate was centrifuged at 1000 rpm for 60 sec. The supernatant was then removed, and the cell pellet was resuspended in a cell isolation solution containing 0.75 mmol/l Ca^2+^. The last step was repeated 4 times. Cells were used within 6-8 hours after the isolation. During this time, the cells retained their brick-like shape with sharp edges and a clear sarcomere patterning.

### Action potential measurement

A drop of cell suspension was added to the bath chamber mounted on the stage of an inverted microscope (IX-71, Olympus, UK) and left undisturbed for 5-10 min to allow the myocytes to adhere to the bottom of the chamber. After that, the cells were superfused at 2 ml/min flow rate with Tyrode’s solution containing (in mM) NaCl 140, KCl 5, CaCl_2_ 1.8, MgCl_2_ 1.0, NaH_2_PO_4_ 0.33, HEPES 10.0, Glucose 10.0; with pH adjusted to 7.4 using 1M NaOH. The perforated-patch current-clamp technique was used for recording the action potentials with 50µM β-Escin in the pipette solution as a perforating agent (26). The patch pipettes were pulled from 1.2 mm borosilicate capillaries (Sutter Instruments, USA) using a two-stage puller (Model PP-830, Narishige, Japan) and fire-polished to facilitate the giga-seal formation. The polished pipettes had a 2 - 2.5 MΩ resistance when filled with the internal pipette solution. The pipette solution for these experiments was composed of the following (in mM): KCl 130, CaCl_2_ 0.5, MgCl_2_ 1.0, Mg-ATP 0.5, EGTA 5.0, and HEPES 10.0. The pH was adjusted to 7.3 with 1M KOH. An appropriate quantity of the 50-mM β-Escin stock solution was added to the pipette solution just before the experiment to achieve a 50-µM final concentration. The presence of β -Escin in the pipette did not interfere with the seal formation. After obtaining a high-resistance (2-5 GΩ) seal, the membrane penetration usually started within 40-60 seconds and a stable access resistance of 25-40 MΩ was obtained within 10-15 min. The cells that failed to achieve this level of access resistance were discarded from further experimentation.

An Axopatch 200B amplifier (Molecular Devices, Sunnyvale, CA, USA) was used for perforated-patch current clamp recording of cardiac action potentials. The resting potential was shifted to -80 mV by applying a negative holding current after recording the existing resting potential (RP) and determining the stimulation threshold by applying 1-millisecond current stimuli of progressively increasing amplitude until an action potential was triggered. The stimulation current amplitude was increased 1.5 times above the threshold and used to elicit the action potentials in all subsequent recordings. The stimuli were generated, and the resulting action potentials recorded using a Digitata 1440A digitizer controlled by Clampex 10.6 data acquisition software (Molecular Devices, Sunnyvale, CA, USA). Four series of 100 stimuli each were applied at 1, 2, 5 and 10 Hz frequencies. Several parameters of the action potentials were extracted for further analysis using Clampfit 10.6. All experiments were performed at room temperature.

### Sodium current recording

The experiments were performed simultaneously with the above current-clamp experiments on a second rig assembled around a Nikon Diaphot 300 microscope (Nikon, USA). Voltage-gated Na^+^ current was recorded from isolated myocytes in whole-cell configuration of the patch clamp technique as previously described (27). We used 4 mM Na^+^ concentration in the bath and the pipette for these experiments. The reduced Na^+^ concentration aided in avoiding “voltage escape” and achieving an adequate voltage clamp of large and fast current (28). Cells were initially superfused with normal Tyrode’s solution as described above. After the seal formation, the bath perfusion was switched to I_Na_ solution of the following composition (in mM): NaCl 4, TEACl 110, CsCl 4, CaCl_2_ 0.1, MgCl_2_ 1.0, HEPES 10.0, Glucose 10.0; with pH adjusted to 7.4 using 1M CsOH. To inhibit currents through the L-type Ca^2+^ channels, 15 µM verapamil was added to this solution. The pipettes were pulled from filamented BF150-86-10 borosilicate glass on a processor-controlled Flaming-Brown type micropipette puller (P-97, Sutter Instrument, USA). The pipette resistance was 1.6-2.0 MΩ when filled with the internal solution containing (in mM): CsCl 120, NaCl 4.0 MgCl_2_ 1.0, Mg-ATP 2, EGTA 5.0, HEPES 10.0. The pH was adjusted to 7.3 with 1M CsOH. The pipettes were positioned and sealed to myocytes using a motorised micromanipulator (PatchStar, Scientifica, UK). A seal resistance exceeding 1 GΩ was obtained by applying a gentle negative pressure. The membrane under the pipette was ruptured by strong suction to gain access to the cell interior. After achieving a whole-cell configuration, the cell was dialyzed with the above solution for at least 5 min before initiating the voltage clamp recording. Access resistance was between 4 and 7 MΩ. Up to 75% of this was electronically compensated to improve the speed and reduce the voltage clamp error. Axopatch 200B amplifier was used for whole-cell patch clamp recording. The voltage clamp protocols were generated and currents recorded using the Digidata 1550 digitizer (Molecular Devices, USA) and Clampex 10.6 software. The current output was filtered at 5kHz using a four-poll Bessel filter of the Axopatch 200B and sampled at 20kHz. The cells were held at -80 mV throughout the experiment. The ‘I_Na_ - voltage’ relationship was measured using 100-ms pulses ranging from -80 to +15 mV after a 100-ms pre-pulse to -120 mV. Cells showing a leak current over 150 pA or signs of ‘voltage escape’ were excluded from further analysis.

### L-type Ca^2+^ current recording

The L-type Ca^2+^ current (I_(Ca, L)_) was recorded as described in our previous paper (25).

In brief, I_(Ca, L)_ was recorded using the above patch clamp setup. The analogue signal was filtered using a four-pole Bessel filter with a bandwidth of 5 kHz and digitised at a sampling rate of 10 kHz. Borosilicate glass patch pipettes were fabricated from filamented BF150-86-10. Electrode resistances in these experiments ranged from 3 to 5 MΩ, and seal resistance from 1 to 5 GΩ. The whole cell bath solution contained the following in mmol/l: 145 NaCl, 2 MgCl_2_, 2 CaCl_2_, 10 HEPES and 10 glucose (pH 7.35). The pipette solution contained the following in mmol/l: 140 CsCl, 2 MgCl_2_, 10 TEA Cl, 10 EGTA, 10 HEPES, 1 MgATP (pH 7.25). More than 75 % of the series resistance was compensated using the Axopatch 200B cell capacitance and series resistance compensation circuitry. Experiments were performed at room temperature. The current-voltage relationship and steady-state inactivation were obtained by applying 1000-ms test pulses in the range of -40 mV to +50 mV from a holding potential of -50 mV. After a brief repolarization to -50 mV, a 120-ms pulse to zero mV was applied to measure steady-state inactivation as shown in Figure 4C. The amplitudes of the currents were normalized to the cell membrane capacitance to obtain current densities (pA/pF).

### Data analysis and statistics

All numerical data were extracted from the recorded traces using Clampfit 10.6 and transferred to IgorPro 9.0 (Wavemetrics, USA) for further analysis. Transmembrane voltages were corrected for 3.6 mV junction potential in I_(Ca, L)_ records and 15 mV in I_Na_ records. IgorPro was also used to create figures and to perform all statistical calculations. More details on particular parameters extraction and analysis are provided in Figure legends where appropriate. The Kruskal-Wallis test was used to treat the data showing non-Gaussian distribution (as presented in Figure 2A-C). For normally distributed data, a one-way ANOVA followed by Tukey *post hoc* analysis was used for means comparisons between the three groups of animals except for data presented in Figure 2D where a one-way repeated measurements ANOVA was employed. The null hypothesis was rejected and the difference was considered statistically significant at p < 0.05. The ‘n’ numbers refer to cell numbers recorded from at least four animals per group.

## Results

### General characteristics of animal groups

The animals were delivered from Charles Rivers at 5 weeks of age and kept at the College of Medicine Animal House for 20-25 weeks before the experiments to allow sufficiently long exposure of the heart to obesity and/or diabetes. After this time, the ZF and ZDF rats exhibited significantly (p<0.05) increased body weight and heart weight compared to ZL rats. Non-fasting blood glucose was significantly elevated in ZDF (384.44±20.28 mg/dl) compared to ZF (140.00±4.58 mg/dl) and ZL (118.25±2.62 mg/dl) rats. Non-fasting blood glucose was not significantly (p>0.05) different between ZF and ZL rats. Heart weight/body weight ratio was significantly (p<0.05) increased in ZF (3.27±0.09 mg/g) compared with (ZL 2.71±0.07 mg/g) and ZDF (2.79±0.05 mg/g) rats. In agreement with a previously published work, the blood plasma insulin level was significantly (p<0.05) higher in ZF rats (12.70±1.15 µg/L) compared to ZL (2.09±0.57 µg/L) and ZDF (2.14±0.29 µg/L) rats.

### Comparison of the action potential parameters in myocytes from ZF, ZDF and ZL rats

In our hands, enzymatic isolation of cardiac myocytes from Zucker rats produced cells with somewhat depolarized RP compared to the normal value measured in adult rat ventricular myocytes. Cells from all groups were capable of generating action potentials with an overshoot in response to brief current pulses of sufficient strength. Typical action potential tracings and a rod-shaped appearance with a clear sarcomere patterning of left ventricular myocytes isolated from ZL, ZF and ZDF groups are shown in Figure 1. The left-hand side panels in Figure 1 show superimposed action potentials elicited from either the pre-existent resting potential or from the membrane potential adjusted to -80 mV by applying a constant current. In all three groups of animals, shifting the resting potential to -80 mV resulted in increased upstroke velocity and shortened the duration of the action potentials evoked by suprathreshold stimulation. Averaged values of the RP, holding current required to shift the RP to -80 mV, threshold potential and action potential duration at 50% and 90% repolarization are presented in Table 1. The values of resting and threshold potentials in individual cells are shown in Figure 2 A and B as “violin” plots, a convenient tool for visualising the data distribution in myocytes from ZL, ZF and ZDF groups (29). The resting potential in myocytes from ZF rats was significantly more negative compared to myocytes from ZL control (p = 0.01525), while the difference between the ZDF and ZL groups was not significant. As one would expect, more depolarized cells should have a lower rate of rise of the action potential due to a larger proportion of sodium channels in the state of steady-state inactivation and lower excitability. Indeed, this was found to be the case. As shown in Figure 2C, the upstroke velocity was the fastest in myocytes from ZF rats (p = 0.04146), while the difference between ZDF and ZL groups was not statistically significant.

**Figure 1.**
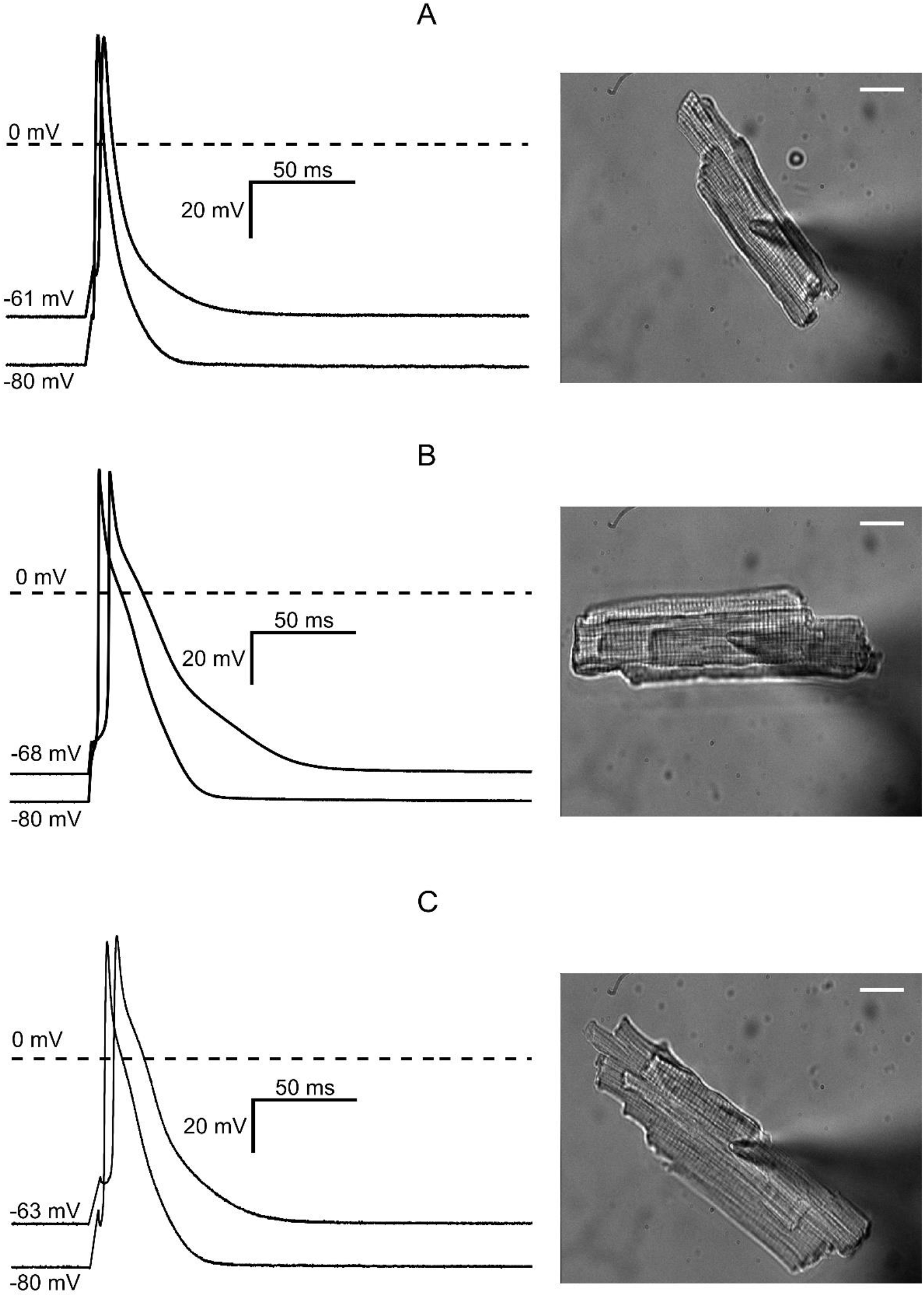
Representative current-clamp recordings from isolated ventricular myocytes. The myocytes shown on the right-hand side of each panel were isolated from A - Zucker Lean (ZL) rats; B - Zucker Fatty (ZF) rats and C - Zucker Diabetic Fatty (ZDF) rats. The left-hand side of each panel shows superimposed action potential tracings evoked by 1-ms threshold current pulse at zero holding current and after applying negative holding current to bring the resting potential to -80 mV. The numbers at the beginning of each trace show the resting potential values before and after the holding current application. The white bars in each image on the right-hand side represent 20 µm.

**Figure 2.**
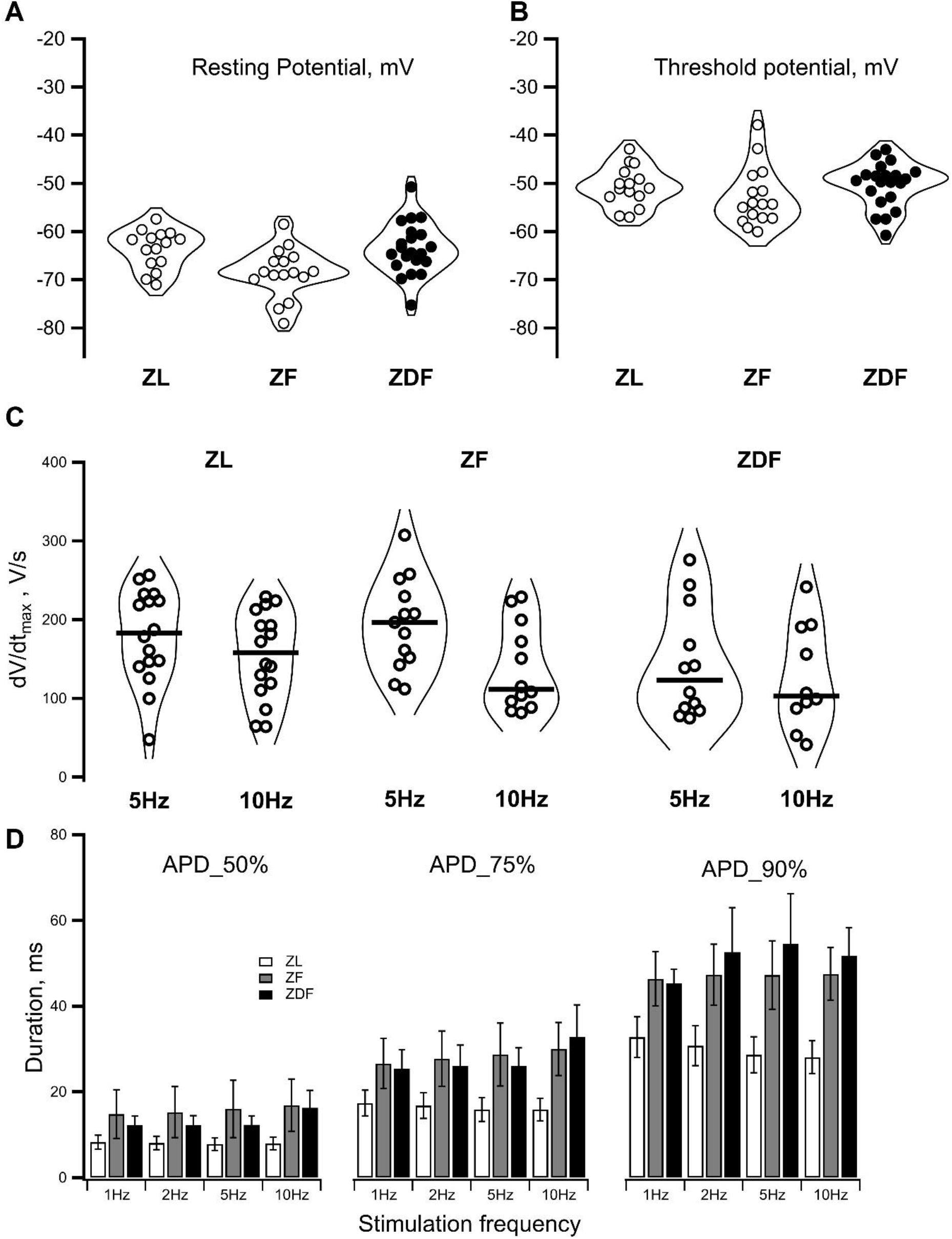
Parameters of resting and action potentials in myocytes from ZL, ZF and ZDF rats. **A** – The violin plot of the pre-existent resting potential values in myocytes from ZL (left), ZF (middle) and ZDF (right) rats. The widest part of each violin corresponds to the highest distribution density of individual data points. ZF rats had more negative resting potential than the other two groups **B** – The violin plots of threshold potentials in ZL, ZF and ZDF rats. Threshold potential was measured as minimal depolarization evoked by a 1-s stimulating current pulse at which an action potential was triggered. Stimuli of lower intensity eliciting only passive electronic responses were considered subthreshold. For all subsequent recordings made in any particular cell, the 1.5 times stronger suprathreshold stimuli were used. **C** – The violin plots of the action potential upstroke velocity (dV/dt _max_) at 5 H and 10 Hz pacing rate in ZL, ZF and ZDF rats. Bold horizontal lines in each violin denote the median of each data set. The largest drop in dV/dt_max_ values in response to the increase in stimulation rate from 5 Hz and 10 Hz was observed in ZF rats. **D** – Ventricular myocytes from ZF and ZDF rats show increased duration of the evoked action potentials. The myocytes were paced at 1, 2, 5 and 10 Hz and action potential duration was measured at 50%, 75% and 90% repolarization. White, grey and black bars represent the mean values correspondingly in ZL, ZF and ZDF rats. The error bars represent the standard deviation.

**Table 1.** Electrophysiological parameters of ventricular myocytes from the ZL, ZF and ZDF rats.

| Action Potential Parameters<br>(Mean ± SEM) | ZL<br>(n=15) * | ZF<br>(n=15) | ZDF<br>(n=13) |
| --- | --- | --- | --- |
| Resting potential (RP) ** | -63.6 ± 1.0 mV | -68.5 ± 1.3 mV | -63.5 ± 1.2 mV |
| Holding current at -80 mV | -0.134 ± 0.048 nA | -0.169 ± 0.022 nA | -0.156 ± 0.051 nA |
| Threshold potential | -50.5 ± 1.0 mV | -53.8 ± 1.5 mV | 50.3 ± 1.0 mV |
| Action potential amplitude | 128.2 ± 3.0 mV | 128 ± 2.0 mV | 128 ± 2.5 mV |
| APD at 50% repolarization | 9.2 ± 1.7 ms | 15.5.3 ± 3.0 ms | 12.2.9 ± 3.4 ms |
| APD at 90% repolarization | 31.7 ± 5.1 ms | 49.0 ± 6.2 ms | 45.0 ± 10.1 ms |

\* The n numbers refer to the number of cells from at least 4 animals per group.
\*\* Measured in each cell before applying the holding current required to set the RP at -80 mV.

Concomitantly, the threshold potential was lowest in the ZF group reflecting their higher excitability (Figure 2B). The upstroke velocity (dV/dt_max_) was dependent on the frequency of stimulation. When paced at 5Hz, which is close to a physiological heart rate in six-month-old Zuker rats (9), dV/dt_max_ was faster than at a pacing rate of 10Hz in all groups of animals (Figure 2C). The biggest drop in dV/dt_max_ was observed in ZF rats: from 193 ± 16.1 V/s at 5 Hz to 136 ± 16.0 V/s at 10 Hz (p< 0.05, n=13). Notably, ZDF rats showed significantly lower dV/dt_max_ at all stimulation frequencies.

### Sodium current in left ventricular myocytes from ZL, ZF and ZDF rats

The low dV/dt_max_ in ZDF rats implies that there could be a diabetes-induced decrease in the expression of sodium channels and/or diabetes-induced changes in their gating. We tested this by comparing the amplitudes and voltage dependence of the inward sodium current in myocytes from ZL, ZF and ZDF rats. Figure 3A shows representative traces of the I_Na_ recorded in myocytes from the lean, fat and fat diabetic animals. The current waveforms showed similar time and voltage dependence (Figure 3A). In all three groups, the current was activated at potentials above -70 mV and the peak of the ‘I_Na_-V_m_’ curve occurred at -40 mV. There were no statistically significant shifts of the ‘G_Na_-V_m_’ curve along the voltage axis between the three groups of animals (Figure 3B, right-hand side panel). The time constant of inactivation and its voltage dependence were similar in all three groups. The only difference detected in these experiments was that the I_Na_ amplitude was significantly higher in ZF rats compared to ZL control (Figure 3B, left-hand side panel). Such augmentation of the I_Na_ was not observed in the ZDF group. The diabetic rats exhibited a diminished I_Na_ compared to the ZF and ZL groups, which would explain the low upstroke velocity observed in diabetic animals. Having established the reason for the diminished upstroke velocity of the ventricular action potential, we next investigated the relation between the action potential duration and frequency of stimulation. In ZL rats, the action potential durations measured at 50%, 75% and 90% repolarization were similar at stimulation frequencies ranging from 1 to 10 Hz.

**Figure 3.**
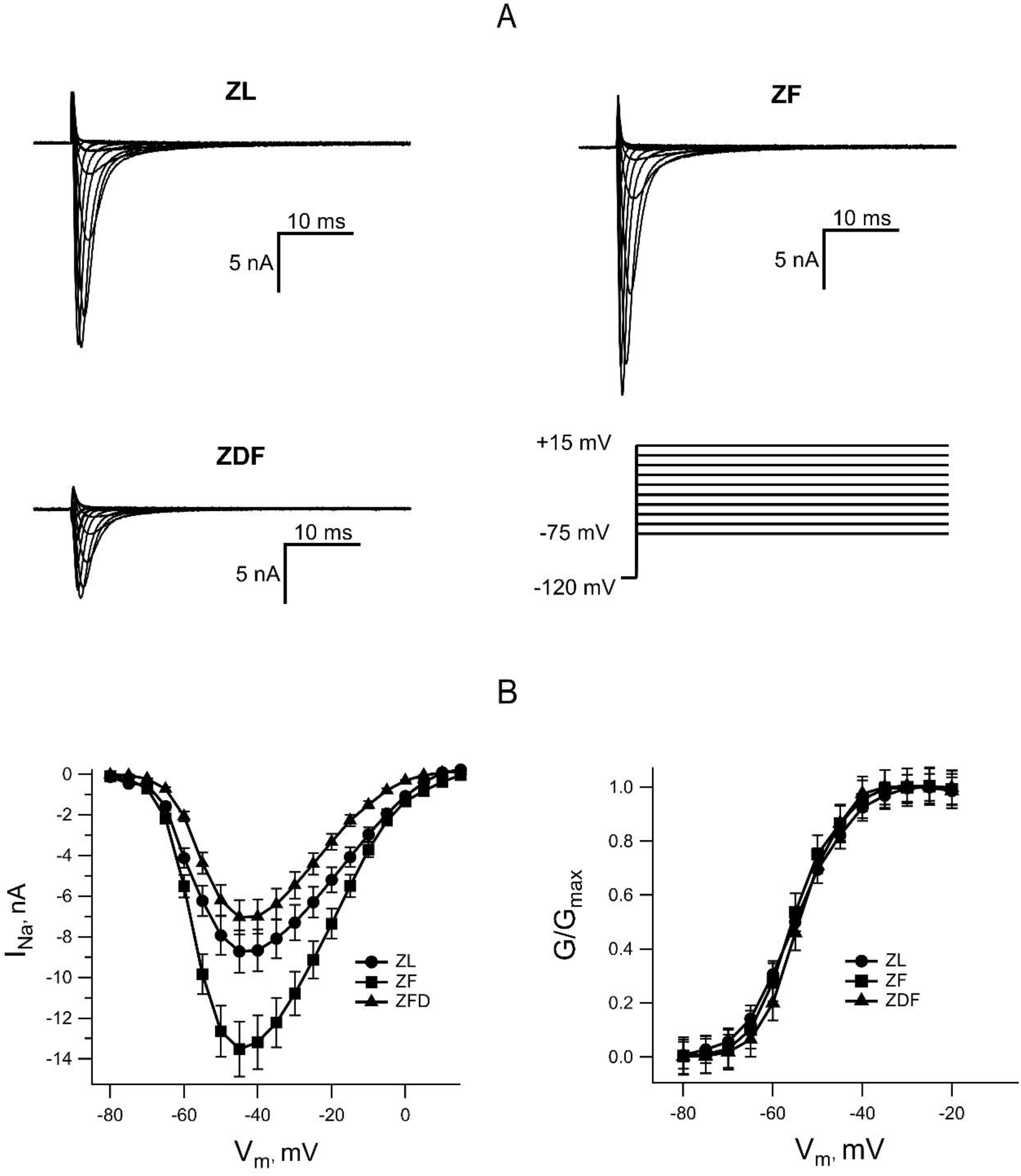
Comparison of voltage-gated Na^+^ currents in myocytes from ZL, ZF and ZDF rats. **A** – Representative traces of I_Na_ elicited by depolarizing voltage clamp pulses (the voltage protocol is shown in the lower panel on the right). **B** - Averaged ‘I_Na_-V_m_’ curves (left-hand side panel) show a significantly increased I_Na_ amplitude in ZF rats in comparison to ZL and ZDF rats. The right-side panel shows the voltage dependence of I_Na_ activation. The sodium conductance (G) was derived from the ‘I_Na_-V_m_’ curves on the left and normalized to the maximum (G_max_) in each group of rats. The peak of the ‘I_Na_-V_m_’ curve was significantly higher in the ZF group. There was no shift in the ‘G_Na_-V_m_’ curves along the voltage axis in any of the three groups of animals.

However, the fatty and diabetic rats showed statistically significant prolongation of their ventricular action potentials (illustrated in Figure 2D). Interestingly, the duration of repolarization was increased at all voltage levels where it was measured i.e., at 50%, 75% and 90% of the action potential amplitude suggesting that the observed elongation was due to the impediment of several ionic currents flowing through the membrane during an action potential.

### L-type Ca^2+^ current in left ventricular myocytes from ZL, ZF and ZDF rats

Multiple studies have described the involvement of several components of voltage-gated K^+^ current as well as voltage-independent inward rectifier K^+^ current in diabetes-induced elongation of ventricular action potential duration. In addition, the contribution from I_(Ca, L)_ “window current” has been suggested although reports on the I_(Ca, L)_ remodelling in diabetic myocardium remain inconsistent. Hence, we have compared the I_(Ca, L)_ densities and voltage dependence of steady-state activation/inactivation parameters in ventricular myocytes from ZL, ZF and ZDF rats. The double-pulse voltage protocol used in this series of experiments allowed the estimation of steady-state activation and inactivation parameters (Figure 4D) simultaneously with the peak amplitude of I_(Ca, L)_ (Figure 4B). Figure 4A illustrates typical current traces elicited by the voltage pulses shown in Figure 4C. During the initial (conditioning) 1-s pulse, the membrane voltage was stepped between -40 and 0 mV in 5 mV increments and between 0 and +50 mV in 10 mv steps. This evoked inward currents that that reached a peak and then inactivated to a steady-state level (see Figure 4A) yielding a current-voltage relationship typical of the voltage-gated L-type Ca^2+^ current. After each 1-s conditioning pulse, the membrane voltage was stepped back to the holding potential for 20 ms and then a 120-ms pulse to 0 mV to measure the fraction of the channels remaining available for activation. The successive conditioning pulses were repeated with 20 s intervals between them, which provided sufficient time for full recovery of Ca^2+^ channels from inactivation during the preceding conditioning pulse. The 5-mV increments in conditioning pulses between -40 and 0 mV allowed for a more precise determination of the I_(Ca, L)_ activation threshold and the position of the overlap between steady-state activation and inactivation curves (“window” current) along the voltage axis. As shown in Figure 4D, there was no significant difference in the “window” current between myocytes from ZL, ZF and ZDF rats. The voltage dependence of the I_(Ca, L)_ was nearly identical between the three groups of animals as judged by Boltzmann’s fit to the steady-state activation and inactivation curves (not illustrated). Likewise, the maximum densities of the L-type Ca^2+^ currents were identical in myocytes from ZL and ZDF rats. There was a trend towards an I_(Ca, L)_ increase in the ZF rats although this did not reach statistical significance (p=0.06375).

**Figure 4.**
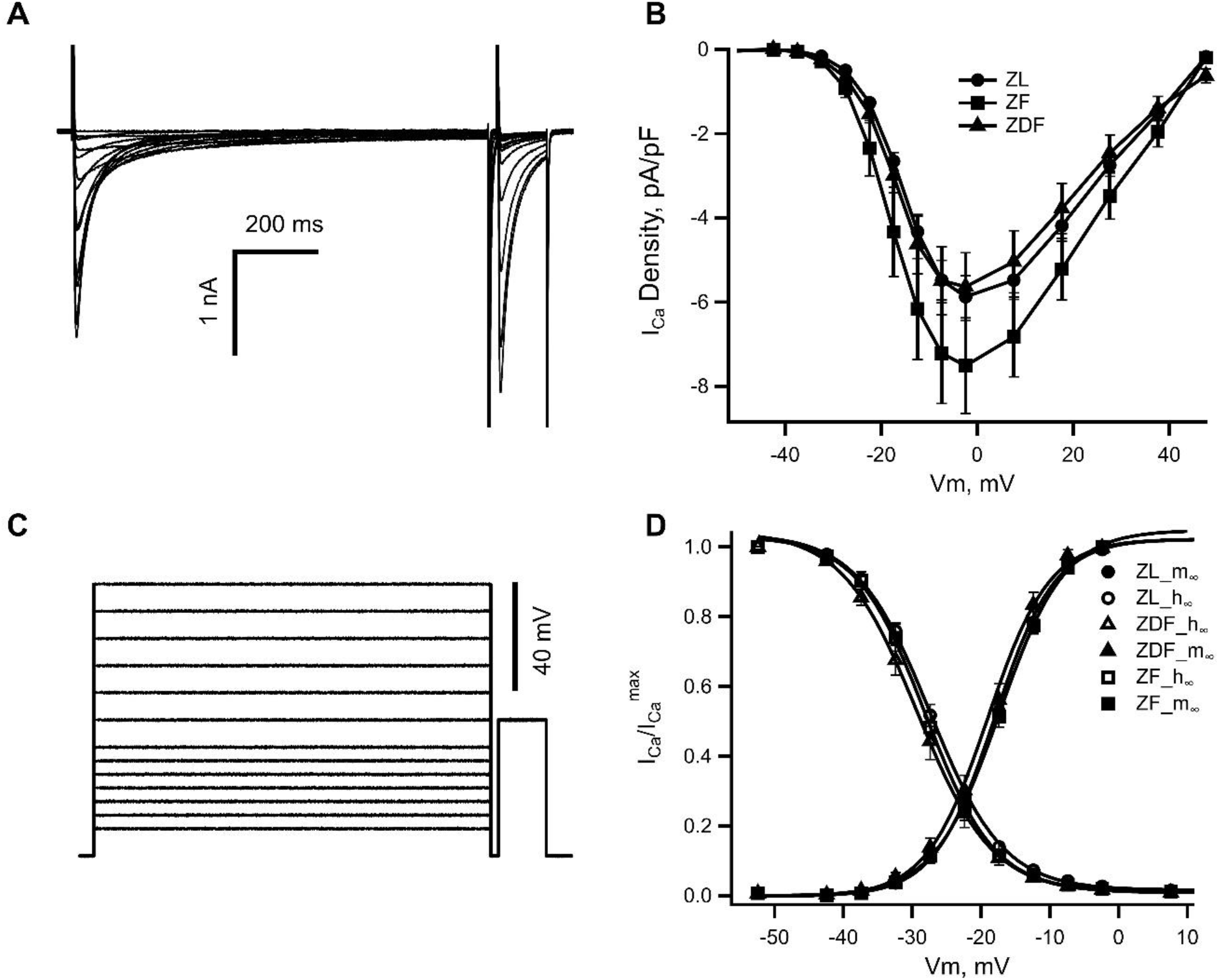
Properties of the L-type Ca^2+^ current (I_Ca, L_) in myocytes from ZL, ZF and ZDF rats. **A** - Typical records of _I(Ca, L)_ from a left ventricular myocyte of a ZF rat. **B** – ‘I_(Ca, L)_ -V_m_’ relationships in ZL, ZF and ZDF rats. Data points represent means ± SEM; n = 15 ZL, 15 ZF and 13 ZDF cells from 4 ZL, 6 ZF and 5 ZDF rats. **C** – an illustration of the voltage clamp protocol used for evaluation of the I_(Ca, L)_ characteristics. **D** – a superposition of steady-state activation and inactivation curves reveals similar voltage dependence of the I_(Ca, L)_ gating in ZL, ZF and ZDF rats.

## Discussion

The Zucker rat model has been one of the most useful small animal models to study the impact of T2DM on the cardiovascular system (5–7,9,30,31). While many metabolomics and biochemistry studies have been published in the literature, there is a paucity of electrophysiological data available (2,9,30,32). In this study, we investigated the remodelling of the left ventricular action potentials in ZL, ZF and ZDF rats. Our results show the prolongation of ventricular action potentials in ZF and ZDF rats at 1-10 Hz pacing frequencies. These results are in agreement with the literature findings (11,17). The pathophysiological significance of this phenomenon is that the long action potential duration may lead to the development of early afterdepolarizations that underlie triggered activity in the ventricle seen on ECG recordings as torsades *des pointes* (33), a specific type of polymorphic ventricular tachycardia that can lead to sudden cardiac death (34,35). Multiple electrophysiological mechanisms have been implicated in the long QT syndrome. A well-established view is that cardiac delayed rectifier voltage-gated and voltage-independent inward rectifier potassium channels play a pivotal role in this phenomenon (34,36,37). More recently the contribution from the L-type Ca^2+^ window current and resurgence of the late Na^+^ current to the development of long action potential and early afterdepolarizations (38–40). We investigated whether the biophysical properties of the L-type Ca^2+^ channels were altered in ZF and ZDF rats compared to control ZL rats and found that this was not the case. In the Zucker Rat model, neither L-type Ca^2+^ channel gating nor I_(Ca, L)_ density are affected by obesity or diabeticity. Our experiments produced no evidence for a non-inactivating sodium current. Furthermore, the I_Na_ inactivation kinetics were similar in all three animal groups ruling out the possibility that the late component of I_Na_ played a role under our experimental conditions. However, we cannot fully reject the possibility of late sodium current appearance under specific hormonal conditions, e.g. in the presence of high levels of angiotensin aldosterone as found in hypertensive diabetic individuals. The use of the Zucker rat model in our study uncovered the differential effects of obesity and T2DM on excitability and the action potential upstroke velocity: a lower threshold and some acceleration of the dV/dt_max_ in obese rats vs. a higher threshold and a noticeable reduction of the dV/dt_max_ in diabetic animals (Figure 2 B & C). The results of our patch-clamp experiments revealed that the observed difference in dV/dt_max_ between ZF and ZDF rats was due to increased I_Na_ amplitude in the fatty non-diabetic group and diminished I_Na_ expression in their diabetic counterparts. This is compatible with the idea that a previously described impairment of electrical conduction in diabetic myocardium (12,19,22) is, in large part, due to the diminished expression of I_Na_ in ventricular myocytes of diabetic animals. Indeed, targeted transgenic disruption of cardiac sodium currents in mice resulted in impaired AV node and ventricular conduction (24). Similar results were obtained in a rabbit model of T1DM (41). Taken together, these findings suggest that dysfunction of myocardial sodium channels is a major cause of impaired electrical conduction.

The differential effects of obesity *per se* and obesity with diabetes on the action potential upstroke velocity suggests that diabetic individuals in contrast to their obese non-diabetic counterparts could be at a higher risk of developing re-entry arrhythmias and conduction block.

In conclusion, the results of our study show a substantial and dissimilar remodelling of the ventricular action potential waveforms in obese non-diabetic and obese diabetic animals underlined by the differential up- and down-regulation of I_Na_ in ventricular myocytes.

## Notes

### Competing Interest Statement

The authors have declared no competing interest.

